# Enhanced early IgG-mediated complement deposition in the development of chronic chikungunya virus disease

**DOI:** 10.64898/2026.08.11.743569

**Authors:** Mrunal Gosavi, Hannah Kamphaugh, Hannah M. Schmidt, Victoria Callahan, Megan M. Dunagan, Jennifer L. Kwan, Liliana Encinales, Alexandra Porras-Ramírez, Alejandro Rico-Mendoza, Aileen Chang, Julie M. Fox

**Author notes:** Corresponding author: Julie M. Fox, Emerging Virus Immunity Unit, Laboratory of Viral Diseases, National Institute of Allergy and Infectious Diseases, National Institutes of Health, Bethesda, MD, 20892. Aileen Chang, Department of Medicine, George Washington University, Washington, DC. **Conflict-of-interest statement:** The authors have declared that no conflict of interest exists.

## Abstract

Chikungunya virus (CHIKV) disease typically resolves following acute infection; however, some individuals develop chronic CHIKV disease (CCD) characterized by persistent, debilitating joint pain. Antibodies help clear CHIKV through neutralization and Fc effector functions. Previous studies have associated CCD with a poor neutralizing antibody response; however, the role of Fc effector functions in CCD development remains unclear. Here, purified IgG from post-acute serum of individuals who either resolved CHIKV disease or developed CCD was evaluated for IgG activity and Fc effector functions to identify correlations with disease progression and biomarkers for CCD. Resolution was associated with higher levels of CHIKV-specific IgG and IgG1 and stronger CHIKV neutralization. Regression analysis identified bulk IgG2 as a strong predictor of CHIKV disease progression. Development of CCD was associated with enhanced IgG-mediated complement deposition on infected cells. These findings suggest that localized complement activation at sites of infection may contribute to persistent inflammation underlying CCD.

## Introduction

Chikungunya virus (CHIKV) is a mosquito-transmitted RNA virus in the alphavirus genus that has caused sporadic, explosive outbreaks worldwide. In late 2013, CHIKV emerged in the Caribbean and quickly spread through Central and South America with autochthonous transmission in Florida and Texas (1, 2). More recently, in 2025, a locally acquired case of CHIKV was reported in New York (3). Since emerging in the Americas, over 3.5 million suspected or confirmed CHIKV cases have been reported in this region (2). During acute infection, individuals can develop fever, rash, myalgia, and severe polyarthritis and polyarthralgia. On average, over 70% of infected individuals still experience severe joint pain three months post-infection, and about a quarter of the infected individuals have chronic arthritis and debilitating joint pain that persists for at least one year following infection (4). Although CHIKV infection is rarely fatal in healthy individuals, significant social and economic loss can occur during and after CHIKV outbreaks (5, 6).

While comorbidities and risk factors, such as age, sex, and acute disease severity, have been correlated with an increased likelihood of developing chronic CHIKV disease (CCD) (7), the cause of CCD remains ill-defined. The presence of persistent CHIKV RNA replication in joint-associated tissue, ultimately promoting inflammation, and the development of an autoimmune response from persistent immune activation have been proposed as potential causes of CCD (8). Previous work has shown persistent CHIKV antigen and/or viral RNA in macrophages, fibroblasts, and myofibers from an individual with CCD and at late time points in animal models (9–11). More recently, a study in mice showed the presence of replicating CHIKV RNA in macrophages in joint-associated tissue at chronic time points (12). However, the detection of persistent CHIKV RNA or antigen in humans at chronic time points is inconsistent (13). Persistent immune activation may be related to other triggers, such as CHIKV peptides and RNA fragments, which could be more challenging to detect. These studies suggest that effective clearance of CHIKV-related immune triggers may limit progression to CCD.

Antibodies are critical for eliminating infectious CHIKV in circulation and are sufficient to prevent lethality in severely immunocompromised mice following CHIKV infection (14, 15). CHIKV-specific antibodies can block infection at various stages of the viral life cycle, including binding, entry, fusion, and egress (16, 17). Additionally, the CHIKV surface glycoproteins (E2 and E1) traffic to the surface of infected cells, allowing antibodies to bind and further engage with immune components, like Fc receptors and the complement component, C1q, to mediate clearance. Numerous mouse and human IgG monoclonal antibodies (mAbs) have been described that are highly protective in vivo (15, 16, 18, 19). Neutralization is generally sufficient to prevent or reduce infection when the mAbs are administered before CHIKV infection (20). However, therapeutic administration of mAbs required Fc-Fc gamma receptor (FcγR) interactions on monocytes for optimal infected cell clearance and disease resolution (21), highlighting the importance of antibody Fc effector functions during CHIKV infection.

Previous studies have shown that a suboptimal antibody response was associated with the progression to CCD (22–24). Neutralizing antibody activity during the febrile stage of CHIKV infection predicted resolution of CHIKV disease after a 20-month follow-up (24). However, it remains unclear if other antibody characteristics, including IgG subclass, binding properties, and Fc effector function activity, contribute to the resolution of CHIKV disease and could be used as an early predictor of disease progression. Here, using serum-purified IgG collected from individuals following acute CHIKV infection, who either resolved CHIKV disease or developed chronic CHIKV arthritis, together with *in vitro* antibody functional assays, we identified early IgG characteristics and effector functions associated with CHIKV disease progression and predictive of CCD development.

## Results

### Increased IgG quantity and neutralization potency in patients who resolved CHIKV arthritis

Serum samples from clinically confirmed cases of CHIKV were obtained from the Chikungunya Arthritis Mechanisms in the Americas (CAMA) study collected in the Atlántico Department of Colombia (25, 26). The serum was confirmed negative for CHIKV RNA and positive for CHIKV-specific IgG and IgM. Post-acute samples from age-and gender-matched patients were divided into those who resolved CHIKV disease (without joint pain; n = 112) and those who developed chronic CHIKV arthritis (with joint pain; n = 112), based on a 20-month follow-up survey (**Figure 1A**) (25, 26). To assess IgG quantity and quality, IgG was purified from each sample, then evaluated for CHIKV-specific IgG levels and CHIKV neutralization potency. Resolved samples had significantly higher levels of CHIKV-specific IgG compared to the chronic samples (p < 0.0001), as shown by the reduced IgG levels required in our CHIKV VLP ELISA to achieve a positive signal (**Figure 1B**). Additionally, individuals who resolved disease showed increased neutralization of a CHIKV strain from the 2014-2016 outbreak in the Americas, with a lower IC_50_ value, compared to those with chronic disease (p < 0.0001) (**Figure 1C**). Analysis of IgG subclasses determined that individuals who resolved disease exhibited significantly higher levels of CHIKV-specific IgG1 (p < 0.0001) and IgG3 (p < 0.001) (**Figure 1D, 1F**); subclasses previously associated with effective antiviral activity in other viruses, including HIV and SARS-CoV-2 (27–29). IgG2 and IgG4 levels did not differ between the groups (**Figure 1E, 1G**).

**Figure 1.**
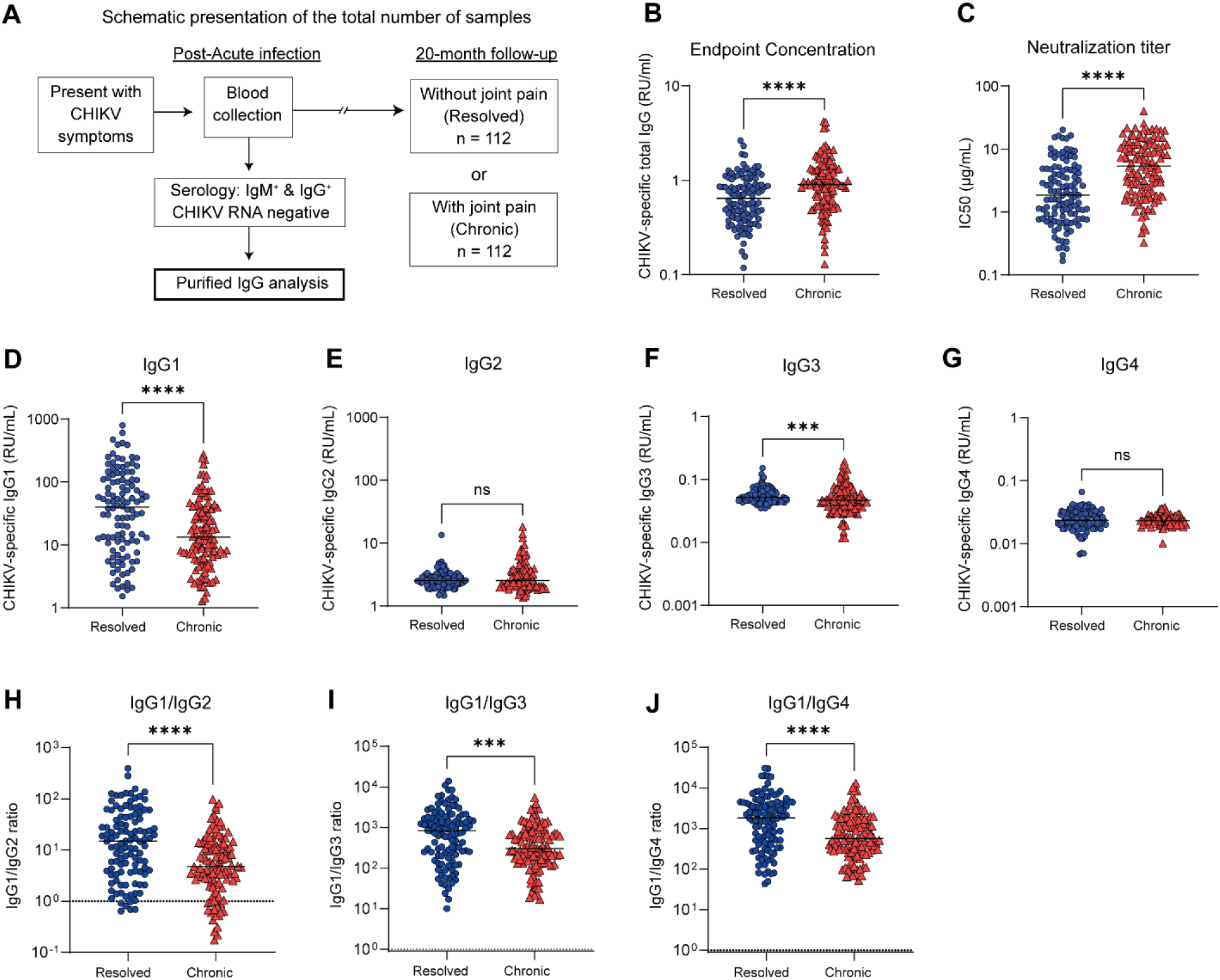
Increased neutralization and IgG response in patients who resolved CHIKV arthritis. (**A**) Schematic presentation of patient cohort and sample collection. IgG was isolated from each serum sample. A total of 121 IgG samples were analyzed per group. (**B**) The endpoint concentration of CHIKV-specific IgG was determined by ELISA. (**C**) CHIKV neutralization was determined by FRNT (IC_50_ value, µg/mL). (**D–G**) CHIKV-specific IgG subclass concentrations for IgG1, IgG2, IgG3, and IgG4 were determined by ELISA compared to a standard curve. The CHIKV-specific IgG subclass concentrations in D-G were used to determine the ratio of (**H**) IgG1/IgG2, (**I**) IgG1/IgG3, and (**J**) IgG1/IgG4. Each dot represents an individual subject; horizontal lines indicate median values. RU stands for relative units. Statistical significance was assessed using a Mann-Whitney test; ****, p < 0.0001; ***, p < 0.001; ns, not significant.

The ratio of IgG subclasses can provide insight into the type and timing of the antibody response. Recovered individuals displayed a higher IgG1/IgG2 ratio (p < 0.0001), suggesting a stronger antiviral response compared to individuals who developed chronic disease (**Figure 1H**). Surprisingly, numerous chronic individuals had more IgG2 than IgG1 (IgG1/IgG2 ratio < 1), indicating a distinct IgG subclass profile associated with chronic disease (**Figure 1H**). While the individuals who resolved CHIKV disease had significantly higher ratios of IgG1/IgG3 and IgG1/IgG4 compared to the patients who developed chronic disease, both groups presented with ratios >1, indicating that the antibody response had matured (IgG1/IgG3 ratio >1) and the environment was more inflammatory than tolerogenic (IgG1/IgG4 ratio > 1) (**Figure 1I-J**).

The samples were down-selected from each group to balance neutralization potency and CHIKV-specific IgG levels for qualitative analysis of functional IgG activities (**Table S1**). The samples were stratified based on neutralization potency into strong neutralizers (IC_50_ < 2 µg/mL; High), moderate neutralizers (IC_50_ = 2–8 µg/mL; Moderate), and weak neutralizers (IC_50_ > 8 µg/mL; Low) or by endpoint IgG dilutions into high (1,600–6,400) and low (400–800) titer groups (**Table S1**). The endpoint dilution was used instead of the endpoint concentration to provide distinct groupings of the samples. The endpoint dilution and endpoint concentration were highly correlated (data not shown), indicating that either representation was acceptable for stratifying the samples. Samples that fit these criteria were randomly selected for the sub-sampling cohort, with age-and gender-balance being maintained (**Table S1**).

Since the resolved samples showed increased CHIKV-specific IgG1 and IgG3, the total IgG subclass levels were quantified in our down-selected samples to determine if the resolved patients naturally skewed toward an IgG1 and IgG3 response or if this response was CHIKV-specific. Bulk IgG1, IgG3, and IgG4 showed no significant differences between patients who either resolved or developed chronic disease (**Figure S1**). Interestingly, bulk IgG2 was significantly elevated in chronic samples (p < 0.0001) (**Figure S1B**). Collectively, these results revealed distinct differences in antibody levels, subclass composition, and neutralization activity between individuals who resolved disease and those who developed chronic CHIKV arthritis, suggesting that qualitative features of the antibody response may be associated with disease outcome.

### Minimal cross-neutralization to other endemic alphaviruses circulating in South American

Previous studies have described cross-reactivity and/or cross-neutralization of CHIKV-immune polyclonal serum and monoclonal antibodies isolated following CHIKV infection to other alphaviruses (17, 19, 30, 31). Mayaro virus (MAYV) and Venezuelan equine encephalitis virus (VEEV) are alphaviruses endemic to South America, with evidence of circulation in Colombia through serological studies (32). With the overlapping geographic distribution and the potential for shared antigenic determinants, we next evaluated neutralization of MAYV and VEEV in our resolved and chronic groups using the down-selected samples. To assess VEEV neutralization, we used a chimeric virus expressing the nonstructural proteins of Sindbis virus (SINV) and the structural proteins of VEEV (SINV-VEEV) (33). Only a few samples neutralized MAYV or SINV-VEEV, with no significant difference observed between the resolved and chronic samples or when the samples were separated based on IC_50_ values against CHIKV [> 8 (Low), 2–8 (Moderate), and < 2 µg/mL (High)] or CHIKV-specific IgG endpoint dilution titers [1600–6400 (High) and 400–800 (Low)] (**Figure S2**). Surprisingly, even though the CHIKV E2 and E1 glycoproteins have higher similarity to MAYV glycoproteins (31), more samples neutralized SINV-VEEV (**Figure S2A** and **D**). These findings suggest that antibodies elicited by CHIKV infection exhibit limited cross-neutralizing activity against MAYV and VEEV, despite the shared phylogenetic lineage.

### Similar IgG avidity and cell surface recognition of CHIKV-infected cells

As the antibody response matures, antibody affinity and avidity increase, resulting in more potent virus neutralization. Since the resolved group had higher neutralization titers against CHIKV, we next evaluated CHIKV-specific IgG avidity in our down-selected sample groups using a chaotropic ELISA with CHIKV virus-like particles (VLPs). The avidity index trended higher in the resolved group compared to the chronic group but did not reach statistical significance (p = 0.07) (**Figure 2A**). Samples from the resolved group with moderate CHIKV neutralization exhibited significantly higher avidity compared to the chronic group (p < 0.05) (**Figure 2B**). No significant differences were observed between the resolved and chronic groups in the low or high neutralization subcategories. When the results were separated based on endpoint dilution titers, no significant differences in the avidity index were observed between resolved and chronic groups (**Figure 2C**). Thus, these results suggest that antibody avidity to the virion is only associated with CHIKV disease resolution in the context of moderate antibody neutralization.

**Figure 2.**
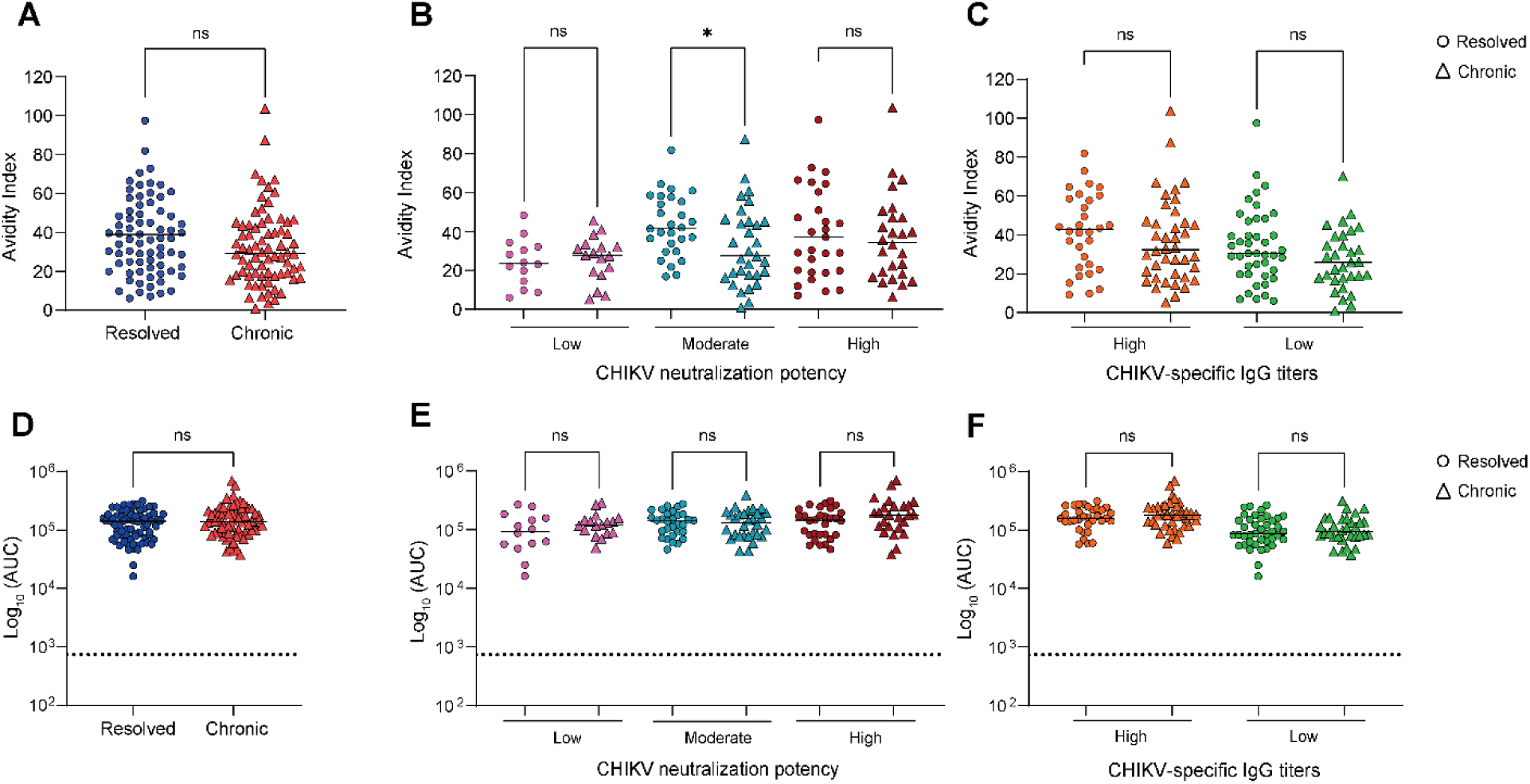
Similar antibody avidity and cell surface binding to CHIKV-infected cells response between resolved and chronic samples. (**A**) The antibody avidity index was determined using a chaotropic ELISA on the purified, down-selected IgG samples (Resolved, n=73; Chronic, n=72). Avidity index was separated by (**B**) CHIKV neutralization (Resolved: low, n=14; moderate, n=29; high, n=30; Chronic: low, n=16; moderate, n=32; high, n=24) or (**C**) CHIKV-specific IgG (Resolved: high, n=32; low, n=41; Chronic: high, n=40; low, n=32) in resolved and chronic samples. (**D**) Binding of the purified IgG to the surface of CHIKV-infected cells was determined by flow cytometry and analyzed by area under the curve (AUC). Subset analysis for cell surface binding based on (**E**) CHIKV neutralization or (**F**) CHIKV-specific IgG between resolved and chronic samples. Each dot represents one individual. Horizontal lines indicate medians. Statistical comparisons were performed using a Mann-Whitney test; significance is indicated as *, p < 0.05; ns, not significant.

During infection, the CHIKV surface glycoproteins are presented on the surface of infected cells (34). Previous studies have shown that antibody engagement with CHIKV antigens on the cell surface can inhibit viral egress and enhance cell clearance through Fc-FcγR interactions (31, 35–38). The previous assays were performed using authentic virus or VLPs; however, antibodies may differentially recognize antigens present on the particle compared to the surface of infected cells. We next assessed IgG binding to the surface of live CHIKV-infected cells by flow cytometry. The area under the curve (AUC) was determined from the integrated median fluorescence intensity (iMFI) of positive cells (**Figure 2D**). Overall, the values were similar between resolved and chronic groups, showing comparable levels of IgG binding to infected cells. Additionally, no significant differences were observed when the samples were stratified by virus neutralization or IgG endpoint dilution (**Figure 2E-F**). These results suggest that resolution of CHIKV disease is not associated with early differences in IgG binding to the surface of CHIKV-infected cells.

### Increased complement-mediated effector function in the chronic group

The Fc region of antibodies can engage with FcγRs on immune cells and C1q to eliminate free virus or virus-infected cells through various effector functions, including antibody-dependent cellular cytotoxicity (ADCC), antibody-dependent cellular phagocytosis (ADCP), and antibody-dependent complement deposition (ADCD) (39). Since failure to clear CHIKV-related immune triggers could contribute to the development of chronic disease, we next investigated antibody Fc-mediated effector functions of the purified IgG from our down-selected samples. First, an ADCC proxy assay was used to assess the interaction between the CHIKV-specific IgG and human FcγRIIIa, which is the dominant FcγR used by NK cells for ADCC in humans (40). In this assay, antibodies are incubated with Raji cells engineered to express the CHIKV structural proteins (Raji-CHIK-VLP), then mixed with Jurkat cells expressing FcγRIIIa (V158 variant). Activation of the effector cells, mediated through FcγRIIIa engagement, results in luciferase activity (**Figure 3A**). No significant difference was observed between the individuals who resolved CHIKV disease and those who developed CCD (**Figure 3B**). When the results were stratified based on CHIKV neutralization or CHIKV-specific IgG titers, we again observed no significant difference in ADCC activity between the resolved and chronic groups (**Figure 3C-D**).

**Figure 3.**
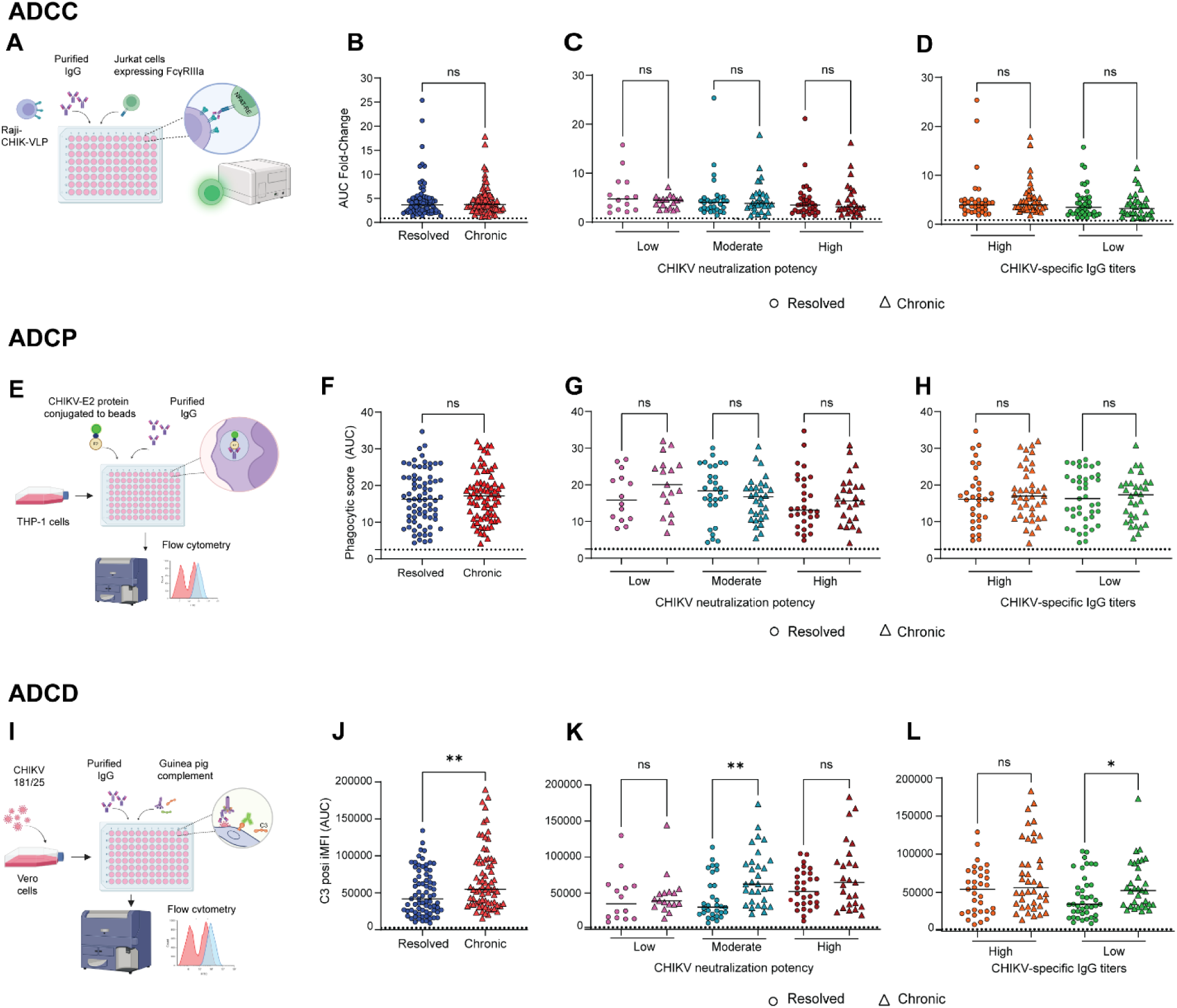
Increased IgG-mediated complement deposition in individuals who developed chronic disease. The purified IgG from the down-selected groups was used to assess Fc-mediated effector functions, including (**A-D**) antibody-dependent cellular cytotoxicity (ADCC), (**E-H**) antibody-dependent cellular phagocytosis (ADCP), and (**I-L**) antibody-dependent complement deposition (ADCD). (**A**) Schematic of the ADCC proxy assay. (**B**) Area under the curve (AUC) analysis was performed using the fold change of ADCC activity compared to the no antibody control. (**E**) Schematic design of the antibody-dependent cellular phagocytosis (ADCP) assay. (**F**) AUC analysis was performed using the phagocytic score. (**I**) Schematic representation of the antibody-dependent complement deposition (ADCD) assay. (**J**) AUC analysis was performed using the integrated median fluorescence intensity (iMFI) of the C3-positive cells. (**C-D**) ADCC, (**G-H**) ADCP, and (**K-L**) ADCD activity were separated by (**C, G**, and **K**) CHIKV neutralization or (**D, H**, and **L**) CHIKV-specific IgG in resolved and chronic samples. The dotted line indicates the average response from 5 healthy human IgG samples. Bars indicate medians. Statistical comparisons were performed using a Mann-Whitney test; significance is indicated as *, p < 0.05; **, p < 0.01; ns, not significant. The images in A, E, and I were created in Biorender.com using a publication license.

Multiple studies have identified antibody-dependent phagocytosis as an important feature of protective anti-CHIKV mAbs to reduce CHIKV infection and disease in mice (31, 36, 41). To investigate ADCP activity, biotinylated CHIKV E2 protein coupled to neutravidin beads was mixed with the purified IgG. The IgG-CHIKV-E2 complexes were then incubated with THP-1 monocyte cells and phagocytic activity was measured by flow cytometry (**Figure 3E**). No significant difference in ADCP activity was observed between the individuals who resolved CHIKV disease compared to those who developed CCD (**Figure 3F**). Further separating the results according to CHIKV neutralization or CHIKV-specific IgG titers demonstrated no significant difference in ADCP activity between resolved and chronic groups (**Figure 3G-H**).

Binding of C1q, in complex with C1r and C1s, to antigen-bound antibodies initiates the classical complement cascade, resulting in the generation of the C3 convertase, which cleaves C3. C3b can then bind to the infected cell surface and trigger the assembly of the membrane attack complex and cell death. Additionally, anaphylatoxins, such as C3a and C5a, can lead to inflammation and chemotaxis, while cells opsonized with C3b and other cleavage products can engage with complement receptors (42). To evaluate IgG-mediated complement activity, we examined the deposition of C3 on cells infected with CHIKV. Purified IgG was incubated with CHIKV-infected cells, followed by the addition of guinea pig complement and detection of C3 on the cell surface by flow cytometry (**Figure 3I**). ADCD responses were significantly elevated in the chronic group compared to the resolved group (p < 0.01) (**Figure 3J**). This increase in C3 deposition was most notable among individuals who developed chronic disease and were categorized as moderate neutralizers or had low IgG titer (**Figure 3K-L**). Together, these findings demonstrate that while cytotoxic and phagocytic antibody functions are similar across disease outcomes, complement-mediated effector activity is selectively elevated early in individuals who develop CCD.

### Increased systemic complement activity in the resolved group

Individuals who progressed to chronic CHIKV disease had an early IgG response that increased deposition of C3 on the surface of infected cells, suggesting enhanced local classical complement pathway activation at the sites of CHIKV replication. To determine whether these differences were reflected systemically, we quantified circulating complement factors associated with the classical pathway, including C1q and C3a, in the serum flow-through following IgG purification. C1q levels did not differ significantly between individuals who resolved or developed CCD, nor when the samples were stratified by CHIKV neutralization potency or CHIKV-specific IgG titers (**Figure 4A-C**). Healthy control samples showed higher C1q concentrations compared to both CHIKV disease groups (**Figure 4A**), suggesting consumption of circulating C1q through activation following CHIKV infection. Surprisingly, C3a levels were significantly elevated in individuals who resolved CHIKV disease compared to those who developed chronic disease (**Figure 4D**). When stratified by CHIKV neutralization potency and CHIKV-specific IgG titers, only resolved samples categorized as low neutralization showed a significant increase in C3a levels (**Figure 4E-F**).

**Figure 4.**
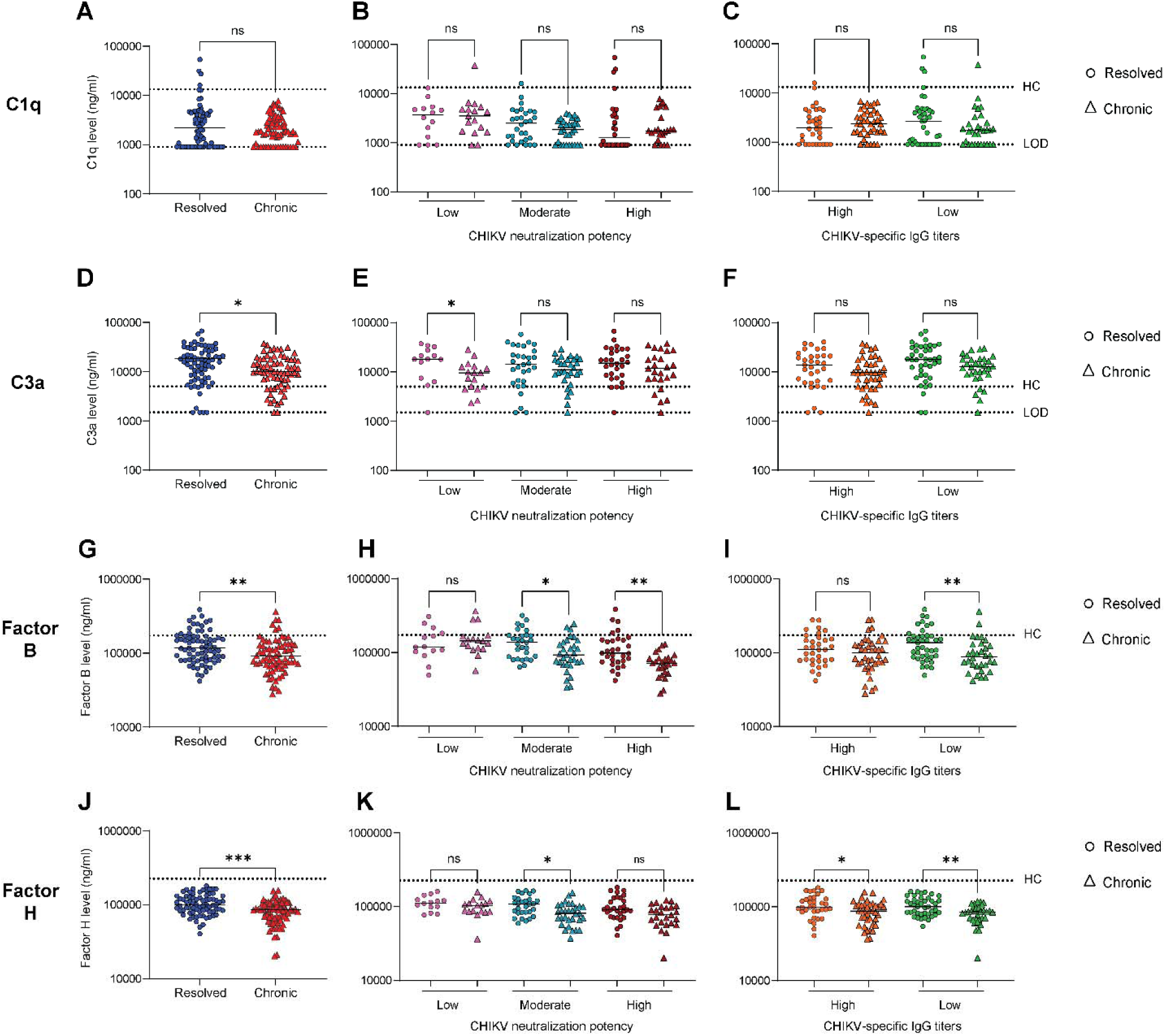
Increased systemic complement activity in individuals who resolved CHIKV disease. After IgG purification, the flow-through fractions from the down-selected samples were analyzed for complement components. **(A-C)** C1q, (**D-F**) C3a, (**G-I**) Factor B, and (**J-L**) Factor H levels were determined by ELISA. The results were stratified by (B, E, H, and K) CHIKV neutralization or (C, F, I, and L) CHIKV-specific IgG. The dotted line labeled healthy control (HC) indicates the average response from 5 healthy human IgG samples. The lower dotted line denotes the limit of detection (LOD) for the assay. Bars indicate medians. Statistical comparisons were performed using a Mann-Whitney test; significance is indicated as *, p < 0.05; **, p < 0.01; ***, p < 0.001; ns, not significant.

Since the three complement activation pathways converge at the cleavage of C3, we quantified complement factors in the alternative (Factor B and Factor H) and lectin (MBL activation) pathways to determine if the increase in C3a in the resolved group was connected to activation of an additional complement pathway (**Figure 4G-L**). Factor B and Factor H were significantly increased in the resolved group compared to the chronic group, suggesting increased alternative pathway activation in individuals who resolved CHIKV disease (**Figure 4G, J**). As with C1q, the healthy controls displayed higher concentrations of Factor B and Factor H than CHIKV-infected individuals (**Figure 4G, J**). When the samples were subcategorized based on CHIKV neutralization potency and CHIKV-specific IgG titers, Factor B was increased in the resolved groups in the moderate and high neutralization and low IgG titer groups (**Figure 4H-I**). For Factor H, individuals who resolved CHIKV disease and had moderate neutralization potency had elevated levels. However, the increase in Factor H was independent of CHIKV-specific IgG titers, as both high and low-titer groups were significantly higher (**Figure 4K-L**). Samples from both CHIKV-infected groups failed to activate the MBL-mediated lectin pathway by ELISA, suggesting that the flow-through contained limited MBL. The MBL likely was consumed following infection, indicating activation of the lectin pathway in both groups. Since there was no difference between the groups in C1q levels, this suggests that increased systemic complement activity and regulation in the resolved group may involve other complement pathway(s) in addition to the classical pathway.

### IgG1 correlates with IgG functionality in resolved individuals, while IgG2 is dominant in chronic individuals

To understand the relationship between IgG characteristics and functionality and disease progression, we performed Pearson correlation separately on the resolved and chronic groups. Since a low value for neutralization (IC_50_ value) and total CHIKV-specific IgG (CHIKV-specific IgG endpoint concentration) indicates better neutralization and more IgG, respectively (**Figure 1B-C**), we used the negative value for the Pearson correlation analysis for ease of interpretation. In patients who resolved CHIKV disease, there was a significant positive correlation of CHIKV-specific IgG1 with total CHIKV-specific IgG, avidity, cell surface binding, ADCD activity, bulk IgG1, and bulk IgG3; while CHIKV-specific IgG2 negatively correlated with neutralization (**Figure 5A**). In contrast, CHIKV-specific IgG1 was negatively correlated with neutralization for the chronic group (**Figure 5B**). Additionally, CHIKV-specific IgG2 was positively correlated with CHIKV-specific IgG3, neutralization, cell surface staining, ADCD activity, and C3a levels and negatively correlated with total CHIKV-specific IgG (**Figure 5B**). These results suggest that patients who resolved CHIKV disease were more dependent on the IgG1 subclass for antibody functionality, while those who developed chronic disease skewed toward the IgG2 subclass. Indeed, when the data were normalized to a Z-score, patients who resolved CHIKV disease developed a coordinated high-quality antibody response characterized by strong neutralization, CHIKV-specific IgG and IgG1, with high avidity and balanced systemic complement response (**Figure 5C**). In contrast, patients who developed chronic disease exhibited increased CHIKV-specific IgG2, Bulk IgG1, IgG2, and IgG3 levels, cell surface binding, and ADCD (**Figure 5D)**. This suggests that chronic patients produce antibodies that bind virus-infected cells and activate the classical complement pathway, but may be less effective at clearing infection, potentially contributing to persistent inflammation.

**Figure 5.**
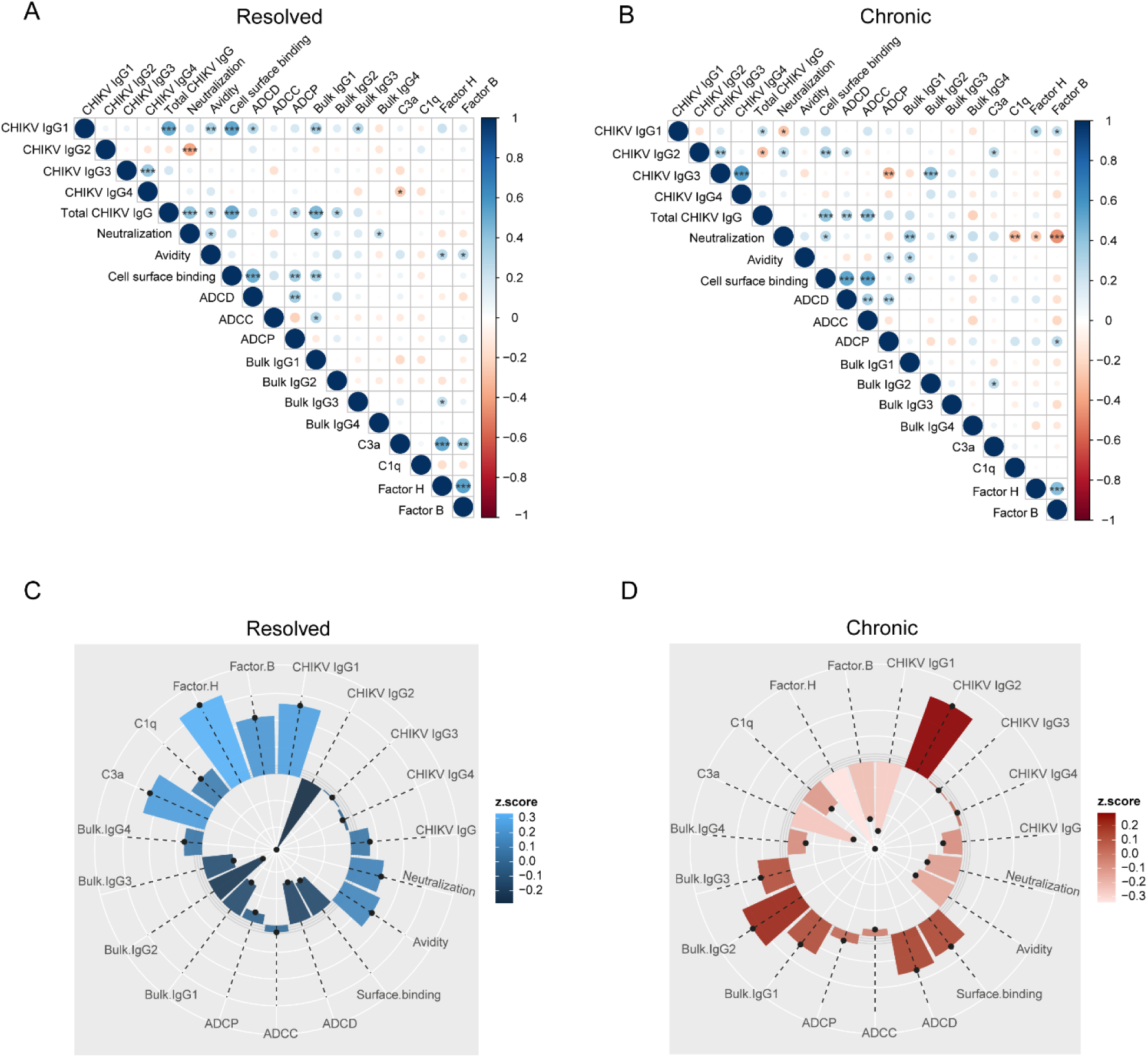
Correlation analysis of IgG responses in resolved and chronic samples. Pearson correlation of the down-selected sample analysis sets in the individuals who (**A**) resolved CHIKV disease or (**B**) developed CCD. The color and size of the circle represent the r value. A stronger correlation has a larger circle. Stars indicate significance (*, p < 0.05; **, p < 0.01; ***, p < 0.001; ****, p < 0.0001). The Z-score for each data point from the analysis was used to generate a petal plot comparing antibody characteristics, Fc effector functions, and circulating complement components for patients who (**C**) resolved CHIKV disease or developed (**D**) chronic disease.

We next performed logistic regression modeling, using the results from each assay as independent variables and the resolved cases as the reference group, with a backward stepwise model variable selection to identify predictors of CHIKV disease. Due to the highly skewed distribution, continuous variables were log_2_ transformed. CHIKV-specific IgG1, IgG3 and IgG4, CHIKV neutralization, cell surface binding, bulk IgG2, and bulk IgG3 levels were identified as significant parameter estimates (**Table 1**). The regression analysis also identified a significant interaction between CHIKV-specific IgG1 and cell surface binding (p = 0.0231). To account for this interaction, the odds ratio (OR) estimates were determined without CHIKV-specific IgG1 included in the model. The most significant predictor for the development of CCD was bulk IgG2 and CHIKV neutralization (p <0.0001), with bulk IgG2 having a higher odds ratio (OR = 4.679) than CHIKV neutralization (OR = 0.755) (**Table 1**). Among all the predictor parameters, the CHIKV-specific IgG4 OR was the highest; however, the wide confidence interval suggests potential outliers altering the distribution. Additionally, the CHIKV-specific IgG4 levels observed were low, indicating this would not be a useful predictor in the clinic. CHIKV-specific IgG3, bulk IgG2, bulk IgG3, and cell surface binding were also significant predictors of CHIKV disease progression. This analysis suggests that quantifying bulk IgG2 in post-acute CHIKV patients could be an early biomarker for potential progression to CCD.

**Table 1.** Predictive modeling for disease progression.

| **Analysis of Maximum Likelihood Estimates** |  |  |  |  |  |
| --- | --- | --- | --- | --- | --- |
| **Parameter** | **DF** | **Estimate** | **Standard Error** | **Wald Chi-Square** | **Pr > ChiSq** |
| Intercept | 1 | -62.2 | 18.13 | 11.8 | 0.0006 |
| CHIKV IgG1 | 1 | 10.3 | 4.87 | 4.47 | 0.0345 |
| CHIKV IgG3 | 1 | -2.35 | 0.840 | 7.80 | 0.0052 |
| CHIKV IgG4 | 1 | 2.52 | 0.931 | 7.32 | 0.0068 |
| Neutralization | 1 | -0.350 | 0.0811 | 18.6 | <.0001 |
| Cell surface binding | 1 | 5.02 | 1.49 | 11.3 | 0.0008 |
| Bulk IgG2 | 1 | 1.92 | 0.362 | 28.2 | <.0001 |
| Bulk IgG3 | 1 | 0.677 | 0.221 | 9.39 | 0.0022 |
| CHIKV IgG1*Cell surface binding | 1 | -0.935 | 0.412 | 5.16 | 0.0231 |

| **Odds Ratio Estimates** |  |  |  |
| --- | --- | --- | --- |
| **Effect** | **Point Estimate** | **95% Wald Confidence Interval** |  |
| CHIKV IgG3 | 0.122 | 0.0320 | 0.461 |
| CHIKV IgG4 | 10.3 | 1.64 | 64.8 |
| Neutralization | 0.755 | 0.659 | 0.865 |
| Bulk IgG2 | 4.68 | 2.69 | 8.15 |
| Bulk IgG3 | 1.97 | 1.34 | 2.89 |
| Cell surface binding | 2.62 | 1.09 | 6.26 |

### Antibody avidity and infected cell binding increase with IgG titers, while antibody avidity and complement deposition increase with neutralization capacity

To examine the overall CHIKV-specific antibody response, we pooled the resolved and chronic down-selected samples and reanalyzed the antibody binding and functional response, then stratified the data according to CHIKV-specific IgG titers and neutralization values (**Figure 6A-J**). This analysis could identify relationships between antibody levels, neutralization potency, and the efficacy of Fc-mediated immune mechanisms. The avidity index increased with higher levels of CHIKV-specific IgG as well as in samples with high neutralizing capacity (**Figure 6A, F**). Additionally, as anticipated, IgG binding to the infected cell surface was higher in samples with more CHIKV-specific IgG, while neutralization potency did not impact recognition of infected cells (**Figure 6B, G**). When examining Fc-mediated functional responses, increased complement deposition was observed in samples with high neutralizing titers but was not dependent on CHIKV-specific IgG titers (**Figure 6C, H**). Samples with more CHIKV-specific IgG showed increased ADCC activity (**Figure 6D**). However, there was no significant difference in ADCC activity when separated by neutralization potency nor ADCP activity when samples were stratified by CHIKV-specific IgG titers or by neutralization potency (**Figure 6E, I-J**). Overall, these findings demonstrate that antibody avidity is associated with the quantity and quality of the antibody response, while binding to infected cells is positively associated with antibody quantity.

**Figure 6.**
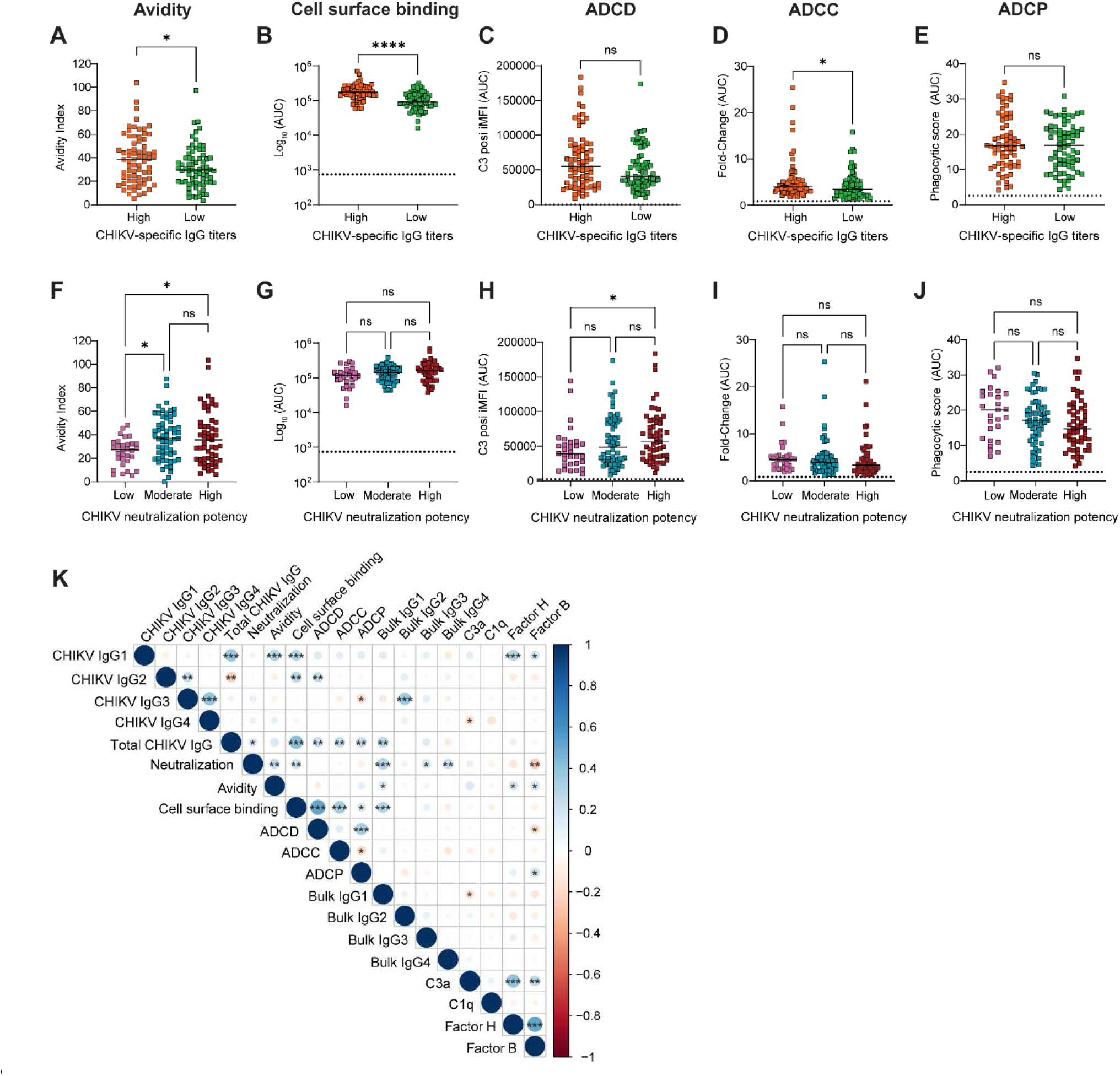
Antibody binding and Fc-mediated immune response to CHIKV infection. The down-selected samples from the resolved and chronic groups were combined, then stratified by (**A-E**) CHIKV-specific IgG or (**F-J**) CHIKV neutralization to evaluate related antibody features. (A and F) Avidity, (B and G) infected cell surface binding, (C and H) ADCD, (D and I) ADCC, and (E and J) ADCP were analyzed. The bars represent the median and the dotted line indicates the response from IgG purified from 5 healthy human IgG samples. Statistical significance was determined using a Mann-Whitney test (A-E) or Kruskal-Wallis with a Dunn’s post-test comparing all groups (F-J) (ns, not significant; *, p < 0.05; ****, p < 0.0001). (**K**) Pearson correlation analysis of the combined down-selected sample set. The color and size of the circle represent the r value. A stronger correlation has a larger circle. Stars indicate significance (*, p < 0.05; **, p < 0.01; ***, p < 0.001 ****, p < 0.0001).

We next performed Pearson correlation on the pooled samples using the negative values for neutralization (IC_50_ values) and total CHIKV-specific IgG (CHIKV-specific IgG endpoint concentration) to assess relationships across the entire data set (**Figure 6K**). The strongest positive correlations were Factor H and Factor B levels (r = 0.53), cell surface binding and ADCD (r = 0.51), total CHIKV-specific IgG and cell surface binding (r = 0.41), and C3a and Factor H levels (r = 0.40). As expected, cell surface binding positively correlated with Fc effector function activity (ADCC, ADCP, and ADCD). The strongest negative correlation was between CHIKV neutralization and Factor B levels (r =-0.25), suggesting a more potent antibody response may reduce activation of the alternative complement pathway.

## Discussion

Our findings identified IgG subclass composition, neutralization potency, and Fc functional profile as key determinants of clinical outcome following CHIKV infection. Individuals who resolved disease exhibited higher levels of CHIKV-specific IgG, increased neutralization capacity, and enrichment of IgG1 and IgG3 subclasses, whereas those who progressed to chronic arthritis demonstrated elevated complement-mediated effector activity. These data suggest that protective immunity is associated with a humoral response that promotes viral inhibition and clearance, while chronic disease may be linked to antibody-driven inflammatory pathways.

Individuals who resolved CHIKV disease had a higher CHIKV-specific IgG1/IgG2 ratio. Surprisingly, numerous individuals in the chronic group had IgG1/IgG2 ratios < 1, indicating more CHIKV-specific IgG2 in circulation than IgG1. IgG1 levels correspond to a stronger antiviral response and effector function activity. In contrast to IgG1, IgG2 has reduced Fc effector function potential, is associated with limited inflammation, and is preferentially generated toward bacterial polysaccharides (28). This increase in CHIKV-specific IgG2 was mirrored in the bulk IgG2 analysis, suggesting that individuals who develop CCD generate a dysregulated immune response or potentially have a secondary bacterial infection, skewing the antibody response (28). While the resolved group showed significantly higher IgG1/IgG3 and IgG1/IgG4 ratios, the ratios for both groups were >1, indicating a mature antibody response consistent with resolution of acute viral infection.

The enhanced IgG1 and IgG3 responses observed in individuals who resolved disease are consistent with the known antiviral properties of these subclasses (43, 44), including their ability to engage Fcγ receptors and mediate effector functions (45). Although FcγR-dependent activities such as ADCC and ADCP did not differ between groups in our assays, the increased representation of IgG1 and IgG3 in resolved individuals potentially contributes to more effective viral clearance through mechanisms not fully captured in vitro. In parallel, the strong association between neutralization potency and disease resolution highlights the importance of antibody quality, in addition to quantity, in limiting viral persistence and downstream pathology.

In contrast, individuals who developed chronic CHIKV arthritis displayed a distinct Fc functional profile characterized by increased complement deposition. Notably, this increase in ADCD occurred with no differences in avidity or antibody binding to infected cells, suggesting a selective skewing of the humoral response or modification of the glycan profile on the Fc region. C1q is known to aid in the clearance of IgG-containing immune complexes through direct binding (46). Complement activation products such as C3a and C5a are potent drivers of inflammation and may contribute to continued joint pathology (47, 48). The association between increased ADCD and lower antibody levels further raises the possibility that, in the absence of efficient viral clearance, complement activation may worsen tissue damage instead of promoting resolution. Although individuals with CCD showed more antibody-dependent complement deposition in vitro, their circulating C3a and Factor B levels were lower compared to the resolved individuals, suggesting reduced systemic complement activation. The increased Factor H levels observed in individuals who resolved disease may also reflect more effective regulation of complement-mediated inflammation, thereby limiting excessive complement activation and tissue damage. This finding suggests that complement activation may be driven by antibody-mediated recognition of CHIKV at sites of virus replication within infected tissue. Localized complement activation may contribute to tissue-specific inflammation with limited changes in circulating complement components.

CCD has been associated with an altered CD4^+^ T cell response (49–51). Previous studies related to autoimmunity have shown that immobilized C1q or C1q bound to apoptotic cells can modulate dendritic cell and macrophage function, leading to suppression of a Th1 and Th17 response in CD4^+^ T cells (52–54). A recent study isolated and restimulated CD4^+^ T cells from individuals with CCD 6 years following infection and showed reduced Th1 subsets and increased expression of TNF-α (50). These findings suggest that antibody-mediated complement activation may also play an indirect role in the establishment of CCD through modification of the CD4^+^ T cell response, although this requires further analysis.

Despite differences in antibody magnitude and function, several features of the humoral response were comparable between groups. Antibody avidity was only modestly increased in resolved individuals and did not consistently distinguish clinical outcomes across stratifications, suggesting that affinity maturation alone is insufficient to explain protection. Similarly, IgG binding to the surface of infected cells was equal between groups, indicating that antigen recognition is preserved regardless of disease condition. Together, these findings suggest that differences in downstream effector functions, rather than antigen binding alone, may be more important in determining disease outcome.

Our findings extend prior studies of CHIKV immunity by highlighting the importance of Fc-dependent antibody functions in shaping clinical progression. While previous work has emphasized neutralizing antibody responses as correlates of protection (24), our data suggest that the balance between protective and potentially pathogenic Fc-mediated activities is also a key factor. This concept is supported by studies in other viral infections, including dengue virus and SARS-CoV-2, where antibody Fc function has been shown to influence both protection and immunopathology (55, 56). In the context of CHIKV, our results suggest that an optimal humoral response may require both effective neutralization and controlled engagement of inflammatory effector pathways.

These findings have implications for vaccines and therapies. Vaccines formulated with TLR agonist adjuvants, such as monophosphoryl lipid A (MPLA) and CpG ODNs, and saponin-based adjuvants, such as Quil-A and Matrix-M, have been shown to elicit a Th1 response in mice characterized by a potent neutralizing antibody response with favorable Fc profiles, particularly IgG2a, which is analogous to human IgG1 (57, 58). CHIKV vaccines administered with these adjuvants may improve protection and reduce the risk of chronic disease. Therapeutic strategies that target complement activation or inflammation may help limit chronic outcomes. In addition, early profiling of antibody functions could help identify individuals at risk of developing chronic arthritis.

This study has several strengths, including the use of a large, gender-matched cohort and a comprehensive evaluation of antibody quantity, quality, and function. However, there are limitations to consider. Samples were obtained at a single post-acute time point, limiting our ability to assess how the antibody response evolves over time. Additionally, IgM is an early, potent neutralizer that primarily mediates effector functions through complement activation rather than classical Fc receptor pathways (56, 59). Future studies should assess IgM responses to better understand early antibody-mediated neutralization, complement activation, and their mechanistic role in CHIKV pathogenesis.

In summary, resolution of CHIKV infection is associated with robust and functionally coordinated antibody responses, whereas chronic disease is linked to increased antibody-dependent complement activity. The absence of elevated systemic complement components suggests that localized, antibody-mediated complement activation at sites of antigen persistence, rather than global complement dysregulation, may contribute to chronic inflammation and disease progression. These findings highlight the importance of antibody functional quality in shaping clinical outcomes.

## Methods

### Study design

As previously reported (25), serum samples from patients with confirmed CHIKV infection were collected in the Atlántico Department, Colombia, as part of an IRB-approved study [George Washington University Committee on Human Research] in January 2015. Samples were serologically confirmed positive for CHIKV-specific IgM and IgG and had no detectable viral RNA. As part of that study, enrolled participants were contacted for a telephone survey 20-months after infection to assess the duration of persistent chikungunya arthritis symptoms (25). Based on the survey results, samples were categorized into resolved (without joint pain) and chronic (with joint pain). We received an equal number (n = 112/ group) of de-identified age-and gender-matched resolved and chronic samples (26).

## Ethics statement

All the participants gave written informed consent to share their samples. The study was approved by the Ethics Committee of the Universidad El Bosque under the protocol “Surveillance of sentinel infectious events prevalent in Colombia” and conducted in accordance with the Declaration of Helsinki (25).

### Cells and viruses

African green monkey cells (Vero; CCL-81), Baby hamster kidney cells (BHK-21; CCL-10), and THP-1 cells were obtained from American Type Culture Collection (ATCC). Vero and BHK-21 cells were cultured in growth media [Dulbecco’s modified essential medium (DMEM) supplemented with 5% heat-inactivated fetal bovine serum (FBS) (HI-FBS, Omega Scientific Inc.) and 10 mM HEPES (Gibco)] at 37°C with 5% CO_2_. THP-1 cells were cultured in maintenance media [RPMI (Thermo Fisher) supplemented with 10% HI-FBS, 1x non-essential amino acids (Gibco), 20 mM HEPES] at 37°C with 5% CO_2._ Expi293 cells were obtained from Thermo Fisher Scientific and were cultured in Expi293 Expression Medium (Gibco) at 37°C with 8% CO_2_ and shaking.

CHIK VLP-Raji cells express the CHIKV structural proteins (strain 37997) under a tetracycline-inducible promoter. The structural polyprotein was amplified from the VLP construct using primers with overhangs to insert an SfiI restriction site at both ends of the PCR product. The PCR product and the pSB-tet-puro vector [generously provided by Dr. Jonathan Yewdell (NIH/NIAID)] were digested with SfiI and ligated at a ratio of 1:3 (vector:insert), then transformed into TOP10 competent cells under carbenicillin selection and sequence confirmed. Raji cells were obtained from ATCC and cultured in RPMI supplemented with 10% HI-FBS and penicillin and streptomycin. The pSBtet-puro-CHIK VLP and the pSB100x [generously provided by Dr. Jonathan Yewdell (NIH/NIAID)] constructs were transfected into the Raji cells using Lipofectamine 3000 (Thermo Fisher). Cells with stable transfection were selected with 5 µg/ml of puromycin (Thermo Fisher). The CHIKV VLP structural proteins were induced with 1 µg/ml doxycycline (Millipore Sigma) in media containing puromycin for 24 hours. CHIKV antigen expression on the surface of live cells was confirmed by flow cytometry.

CHIKV strain 99659 and strain 181/25 were generated from a cDNA clone as described (60) and passaged on BHK-21 cells. Experiments with the CHIKV strain 99659 were performed under BSL3 conditions. MAYV (strain BeH407) was obtained from the World Reference Center for Emerging Viruses and Arboviruses and SINV-VEEV (strain TrD) was generously provided by Dr. William Klimstra (University of Pittsburgh). MAYV and SINV-VEEV were passaged on Vero cells. Virus stocks were titrated by focus-forming assay.

### Purification of human IgG

Antibodies were purified using Protein G HP SpinTrap^TM^ columns (Cytiva) and Ab buffer kit (Cytiva), according to the manufacturer’s instructions with minor modifications. Briefly, the column storage solution was removed by centrifugation. The column was equilibrated with binding buffer. The column was securely capped and serum sample was added and incubated for 4 min with gentle mixing to allow binding. The column was centrifuged to remove unbound material. The serum flow-through was collected and stored at-80 °C. The column was washed twice with binding buffer. For elution, elution buffer was added to the column and mixed by inversion. The column was then placed into a microcentrifuge tube containing neutralizing buffer and centrifuged to collect the eluate. A second elution was performed using the same conditions in a fresh tube containing neutralizing buffer. Eluted fractions were pooled, antibody concentration was checked by NanoDrop (Thermo Fisher), and stored at-20°C.

### CHIKV focus reduction neutralization test (FRNT)

Vero cells were seeded in 96-well plates (2.5 × 10L cells/well) and incubated in CO_2_ incubator overnight at 37°C with 5% CO_2_. Purified human IgG was serially diluted three-fold in infection medium starting at 1:40, mixed 1:1 with CHIKV, then incubated for 1 h at 37°C with 5% CO_2_. Antibody–virus mixtures containing 100 focus-forming units (FFUs) of CHIKV were added to Vero cell monolayers for 1.5 h at 37°C with 5% CO_2_. Overlay media [Minimum Essential Medium (Sigma-Aldrich) with 1% methylcellulose (Sigma-Aldrich), supplemented with 2% HI-FBS, 1M L-glutamine, 1M HEPES, 1% pen/strep, 0.42% sodium bicarbonate (Sigma-Aldrich)] was added onto the cells and incubated for 18-20 h at 37°C with 5% CO_2_. Cells were fixed with 1% paraformaldehyde (PFA; Electron Microscopy Sciences) diluted in PBS for 1 h at room temperature (RT), washed thrice with PBS, and permeabilized with PBS containing 0.1% saponin (Sigma-Aldrich) and 0.1% bovine serum albumin (BSA; Sigma-Aldrich) (perm wash). Viral foci were stained using a mouse anti-CHIKV E2 monoclonal antibody (CHK-11; 500 ng/mL). After incubating for 2 h at RT, cells were washed with wash buffer (PBS + 0.05% Tween-20), followed by an HRP-conjugated anti-mouse IgG (H+L) secondary antibody (1:2000; KPL) for 2 h at RT, then washed with wash buffer. Foci were developed using Tru-Blue substrate (KPL) and counted using a BioSpot reader (ImmunoSpot by CTL). The percent relative infection was calculated relative to virus-only controls. The IC_50_ value was determined using non-linear regression with the top and bottom constrained to 100 and 0, respectively.

### Cross-neutralization assays for other alphaviruses

Vero cells were seeded in a 96-well plate (2.5 x 10^4^ cells/well) and incubated overnight at 37°C with 5% CO_2_. Purified human IgG samples and positive control antibodies [human anti-CHIKV (CHKV-24; Leinco Technologies) and mouse anti-VEEV (VEEV-57; Bio X Cell)] were diluted to 20 µg/mL in infection media, mixed 1:1 with 200 FFUs of MAYV or SINV-VEEV, then incubated for 1 h at 37°C with 5% CO_2_. The antibody-virus mixture containing 100 FFUs of virus was added to the Vero cell monolayer and incubated for 1.5 h at 37°C with 5% CO_2_. Overlay media was added and incubated for an additional 16 h (MAYV) or 20 h (SINV-VEEV) at 37°C with 5% CO_2_. Cells were fixed with 1% PFA diluted in PBS for 1 h at RT, washed with PBS, then permeabilized with perm wash. Foci were stained with CHK-48 (MAYV) or VEEV-57 (SINV-VEEV) (500 ng/mL) overnight at 4°C, washed, then incubated with HRP-conjugated anti-mouse IgG (H+L) secondary antibody (1:2000, KPL) for 2 h at RT. Foci were developed using Tru-Blue substrate and counted using a BioSpot reader. The percent neutralization was calculated relative to virus-only controls.

### CHIK-VLP preparation and purification

Expi293 cells were transfected with CHIKV-VLP plasmid [strain 37997; pCHIK-37997ic; generously provided by Dr. Michael Diamond (Washington University School of Medicine)] using the Gibco™ ExpiFectamine™ 293 Transfection Kit (Thermo Fisher), following the manufacturer’s instructions, and incubated at 37°C with 8% CO_2_ with shaking. The following day, ExpiFectamine™ Transfection Enhancer 1 and ExpiFectamine™ Transfection Enhancer 2 were added to the flask and incubated at 37°C with 8% CO_2_ with shaking for 4 days.

Supernatant was then collected and clarified by centrifugation and filtration. VLPs were pelleted through a 20% sucrose cushion by ultracentrifugation for 4 h at 24,000 rpm and 4°C using a SW32 rotor. The VLP pellet was resuspended in Tris-NaCl-EDTA (TNE) buffer. VLPs were purified through a 20% and 60% discontinuous sucrose gradient by ultracentrifugation for 4 h at 24,000 rpm and 4°C using a SW41 rotor. The visible VLP band at the interface of the 20% and 60% discontinuous sucrose gradient was collected and verified by protein gel electrophoresis. Purified VLPs were buffer exchanged into 1X PBS using the Pur-A-Lyzer™ Maxi Dialysis Kit (Millipore Sigma).

### CHIKV-specific total IgG ELISA

CHIKV-VLP (1 µg/mL) was coated overnight at 4°C on MaxiSorp Thermo Scientifc™ Immuno ELISA plates in coating buffer (carbonate-bicarbonate buffer, pH 9.2). ELISA plates were washed three times with wash buffer and blocked with blocking buffer (PBS + 5% BSA) for 1 h at 37°C. Following blocking, purified human IgG were serially diluted two-fold starting at 1:50 in blocking buffer and incubated for 1 h at RT. Plates were washed three times with ELISA wash buffer and incubated with mouse Anti-Human IgG Fc-HRP (1:5000; Southern Biotech) diluted in blocking buffer for 1 h at RT. Plates were washed three times with wash buffer and developed for 2-5 minutes at RT using TMB One-Step Substrate (Thermo Scientific). The reaction was neutralized using 2M H_2_SO_4_ and absorbance was measured at 450 nm using a BioTek Synergy H1 microplate reader (Agilent). The cut-off value was defined by the OD value for no IgG control plus three times the standard deviation. The endpoint concentration was determined by nonlinear regression of the ELISA OD values across the serial dilution of purified IgG.

### CHIKV-specific IgG subclass ELISA

CHIKV-VLP (1 µg/mL) was coated overnight at 4°C on MaxiSorp Thermo Scientific™ Immuno ELISA plates in coating buffer. ELISA plates were washed three times with wash buffer and blocked with blocking buffer for 1 h at 37°C. Purified human IgG samples were serially diluted two-fold across the ELISA plate starting at 1:50 in blocking buffer. For subclass standards, subclass-specific antibodies [IgG1, human IgG1 anti-CHIKV (CHKV-24; Leinco Technologies); IgG2, human IgG2 isotype control (BioXCell); IgG3, human IgG3 Lambda (Southern Biotech); IgG4, human IgG4 anti-hen egg lysozyme (S228P) (BioXCell)] were diluted to 10 µg/ml (IgG1 & IgG2) or 0.5 µg/ml (IgG3 & IgG4) in blocking buffer then serially diluted two-fold. Plates were incubated for 1 h at RT, then washed three times with wash buffer. Next, plates were incubated with corresponding HRP-conjugated anti-human IgG subclass-specific antibodies [IgG1, mouse anti-Human IgG1 Fc-HRP (Southern Biotech); IgG2, mouse anti-Human IgG2 Fc-HRP (Southern Biotech); IgG3, mouse anti-human IgG3 hinge-HRP (Southern Biotech); IgG4, mouse anti-human IgG4 pFc’-HRP (Southern Biotech)] at a dilution of 1:5000 in blocking buffer for 1 h at RT. Plates were washed three times with wash buffer and developed for 2-5 minutes at RT using TMB One-Step Substrate. The reaction was neutralized using 2M H_2_SO_4_ and absorbance was measured at 450 nm using a BioTek Synergy H1 microplate reader. Standard curves were generated by plotting log_10_[antibody standard] against the experimentally obtained OD value, then used to calculate the concentration of each IgG subclass.

### Bulk IgG subclass ELISA

Goat anti-human IgG cross-adsorbed to mouse serum proteins (Southern Biotech) (1 µg/ml) was coated in coating buffer on Maxisorp immunocapture ELISA plates (Thermo Fisher Scientific) and incubated overnight at 4°C. Plates were washed three times using wash buffer. Plates were blocked at 37°C for 1 h in blocking buffer. Purified human samples were diluted (starting at 1:200 for IgG1 and IgG3 or 1:100 for IgG2 and IgG4) in blocking buffer, then three-fold serial dilutions were performed on the ELISA plates and incubated for 1 h at RT. Plates were washed three times using wash buffer, and incubated with the corresponding IgG subclass HRP-conjugated secondary antibody, as described above, for 1 h at RT. After washing the plates with wash buffer, the plates were developed with TMB One-Step Substrate for 2-5 min at RT. The reaction was neutralized using 2M H_2_SO_4_ and absorbance was measured at 450 nm using a BioTek Synergy H1 microplate reader. Standards were performed as described above for the CHIKV-specific IgG subclass ELISA and used to quantify the level of each IgG subclass in each sample.

### Avidity ELISA

CHIKV VLPs (Strain 37997) (2 µg/ml) were absorbed overnight at 4°C on Maxisorp immunocapture ELISA plates in coating buffer. Wells were washed with wash buffer and blocked with 2% BSA blocking buffer [PBS + 2% BSA (Sigma)] for 2 h at 37°C. Purified human IgG was diluted in 2% BSA blocking buffer to 10 µg/mL and then added to wells for 2 h at RT. Plates were washed with wash buffer, then incubated for 20 minutes at RT with 5M urea (Sigma-Aldrich) or PBS. Plates were then washed three times with wash buffer and reblocked with 2% BSA blocking buffer for 1 h at 37°C before incubation with HRP-conjugated mouse anti-human IgG Fc (Southern Biotech). Plates were washed with wash buffer and developed using TMB One-Step Substrate solution (Thermo Fisher). The reaction was stopped with 2M H_2_SO_4_ and absorbance was measured at 450nm using a BioTek Synergy H1 microplate reader. Relative avidity index (RAI) was calculated for each sample by dividing the OD at 450nm in urea-treated wells by that in untreated wells (PBS) for each individual sample.

### Cell surface staining of CHIKV-infected cells

Vero cells were seeded at 4 x 10^6^ cells in a T-75 flask in cell growth media and incubated overnight at 37°C and 5% CO_2_. The next day, Vero cells were infected with CHIKV 181/25 at a multiplicity of infection (MOI) of 1 in a low volume of infection media (DMEM + 2% HI-FBS + 1:100 HEPES + 1:100 penicillin/streptomycin) for 1 h at 37°C with 5% CO_2_. After 1 h, additional media was added and infected cells were incubated for 18 h at 37°C and 5% CO_2_. Infected cells were rinsed with PBS and trypsinized from the flask. Cells were pelleted then washed with FACS buffer (PBS + 1% HI-FBS + 1mM EDTA). Cells were resuspended in FACS buffer and 5 x 10^4^ cells were seeded into a 96-well U-bottom plate. Following pelleting, cells were stained with three-fold serial dilutions of purified human IgG samples, positive control (CHKV-24; Leinco Technologies), or negative control (healthy human IgG) starting at 10 µg/mL for 1 h at 4°C. Cells were washed in FACS buffer, then incubated with goat anti-human IgG-AF647 (1:2000; Southern Biotech Cat No. 2040-31) in FACS buffer for 1 h at 4°C protected from light. Cells were washed in FACS buffer and fixed in 4% PFA in PBS for 10 minutes at 4°C. Following fixation, cells were washed twice in FACS buffer then resuspended in FACS buffer for analysis. Samples were run on a BD LSRFortessa (Waters Biosciences) and analyzed using FlowJo (Waters Biosciences; version 10.10.0). Voltages were set based on the positive control to ensure reproducibility across assays. The integrated median fluorescence intensity (iMFI) was calculated by multiplying the median fluorescence intensity (MFI) by the percentage of positive cells for each well and the area under the curve (AUC) was determined using Graphpad Prism (Version 10.4.1).

### Antibody-dependent cellular cytotoxicity (ADCC) assay

Purified human IgG, positive control (CHKV-24), and negative control (healthy human IgG) were assessed for ADCC activity using the ADCC reporter bioassay, FcγRIIIa V variant (Promega). Samples were diluted in ADCC assay buffer (RPMI 1640 + 4% low IgG serum) to 10 µg/ml followed by three-fold serial dilutions. Raji-CHIK-VLP cells were thawed and, immediately, 2 x 10^4^ cells /well were seeded in 96-well white-walled, flat-bottom plate (Corning). Serially diluted human IgG was added to the Raji-CHIK-VLP cells and incubated for 1 h at 37°C with 5% CO_2._ Then, 1 x 10^5^ effector cells (hFcyRIIIa V158 Jurkat/NFAT-luc; Promega) were added to each well containing Ab-Raji-CHIK-VLP complex at an effector:target cell ratio of 5:1 and incubated for 6 h at 37°C with 5% CO_2._ Plates were brought to RT for 15 min before the 6 h incubation ended. Reconstituted Bio-Glo Luciferase Assay Reagent (Promega) was added to each well and incubated for 10 min at RT. Plates were read at a gain of 200 Lum on a BioTek Synergy H1 microplate reader, and the luminescence was measured in relative light unit (RLU). The positive and negative controls were included on each plate to ensure reproducibility across assays. The background RLU was subtracted, then divided by the average of the no-antibody control wells to obtain the fold induction. The fold change at each concentration was used to calculate the AUC using GraphPad Prism.

### Antibody-dependent complement deposition (ADCD) assay

Vero cells were seeded at 4 x 10^6^ cells in a T-75 flask in cell growth media. The next day, the cells were washed with PBS and infected with CHIKV 181/25 at an MOI of 1 in infection media for 1 h at 37°C with 5% CO_2_. The cells were rinsed twice then fresh infection media was added. The cells were incubated for 18 h at 37°C with 5% CO_2_. Purified human IgG, positive control (CHKV-24), and negative control (healthy human IgG) were diluted in RPMI + 10% HI-FBS starting at 10 µg/ml with three-fold serial dilutions. Infected Vero cells were trypsinized, pelleted at 1500 x rpm for 5 min at 21°C, then resuspended in RPMI + 10% HI-FBS. Cells were seeded at 5 x10^4^ on the serially diluted antibody and incubated for 1 h at 37°C with 5% CO_2_. Cells were washed twice with RPMI + 10% HI-FBS. Lyophilized low-tox guinea pig complement (Cedarlane) was reconstituted in cold sterile water and diluted 1:50 in RPMI + 10% HI-FBS. Diluted complement (200 µl) was added and incubated for 15 min at 37°C with 5% CO_2_. Cells were washed with quenching buffer (PBS + 1% HI-FBS + 15 mM EDTA), then with FACS buffer (PBS + 1% HI-FBS + 1 mM EDTA). Cells were resuspended in goat anti-guinea pig complement C3-FITC (1:100, MP Biomedicals) and LIVE/DEAD Fixable Aqua Stain (1:500, Invitrogen) in FACS buffer for 30 min at 4°C. Cells were washed twice with FACS buffer, fixed with 4% PFA diluted in PBS, and incubated for 10 min at 4°C. Cells were washed with FACS buffer and resuspended in PBS. Cells were run on a BD LSRFortessa followed by analysis using FlowJo (version 10.10.0). The voltages were set based on the positive control to ensure reproducibility across assays. The percentage of C3^+^ cells was determined on single, viable cells. The iMFI was calculated by multiplying the MFI by the percentage of positive cells for each well. AUC was calculated for the iMFI using GraphPad Prism.

### Antibody-dependent cellular phagocytosis (ADCP) assay

Recombinant CHIKV E2 (Leinco) was biotinylated and desalted. Biotinylated E2 was then conjugated to FluoSpheres NeutrAvidin-labeled microspheres (Thermo Fisher) for 1 h at RT, with vortexing every 10 min. The optimal quantity of biotinylated E2 and beads was determined experimentally. CHIKV E2-coated beads were incubated with three-fold serial dilutions of purified human sample IgG in maintenance media for 2 h at 37°C. THP-1 cells were incubated with CellTrace violet (Thermo Fisher) for 20 mins at 37°C. The THP-1 cells were then added to the immune complexes at a concentration of 4.9 x10^4^ cells/well and incubated 16 h at 37°C in a low-adherence 96-well plate. Cells were then pelleted and washed twice with FACS buffer before being fixed with 4% PFA in PBS and incubated for 10 min at RT. Cells were run on a BD LSRFortessa followed by analysis using FlowJo (version 10.10.0) analysis software. Voltages were set based on the positive control to ensure reproducibility across assays. The phagocytic score was determined by multiplying the MFI by the percent positive cells and further dividing it by 100. The normalized phagocytic score was calculated by subtracting the phagocytic score of a no-antibody control from the phagocytic score of a sample. AUC was calculated for the phagocytic scores using GraphPad Prism.

### Complement ELISAs

Flow-through from each sample after IgG purification was stored at-80 °C until analysis. Factor B was quantified in the flow-through (1:8000 dilution) using the Human Factor B ELISA Kit (Abcam), following the manufacturer’s instructions. Human Factor H, C1q, and C3a were measured using the Human Complement Factor H ELISA Kit (Abcam), Human C1q ELISA Kit (Invitrogen), and Human Complement C3a ELISA Kit (Invitrogen), respectively. The assays were performed according to the protocols and dilutions recommended by the indicated manufacturer’s instructions.

## Statistical analysis

The total number of participants and group-specific sample sizes for the down-selected samples are summarized in Table S1, any change in the sample number is indicated in the respective figure legends. Statistical analysis was conducted using GraphPad Prism (Version 10.4.1). Data are presented as median values. The specific statistical tests and multiple-comparison post-tests, when applicable, are described in the figure legends. For comparisons between two independent groups, the Mann-Whitney *U* test was used. Comparisons among three groups were performed using the Kruskal-Wallis test followed by Dunn’s multiple-comparison test. All statistical tests were two-sided, and a *P* value less than 0.05 was considered statistically significant. Correlation plots were generated in R (version 2026.06, “Mariposa Orchid”) with the corrplot, dplyr, Hmisc, and tidyr packages. Regression analyses were conducted using SAS version 9.4 (SAS Institute Inc., Cary, NC, USA).

## Data availability

All data underlying each figure are available in S1 Data. Additional requests should be directed to the corresponding author. Reagents are available under a material transfer agreement.

## Author contributions

M.G., A.C., and J.M.F. conceptualized the study. M.G., H.K., H.M.S., V.C., M.M.D., and J.M.F. developed methodology. M.G. performed purification of human IgG, CHIKV FRNTs, quantitative bulk IgG ELISA, ADCD assay, and complement component ELISAs. H.K. performed MAYV and SINV-VEEV FRNTs, CHIK-VLP preparation, CHIKV-specific IgG and subclass ELISAs, and cell surface binding assays. H.M.S. performed ADCP assays. V.C. performed ADCC assays. M.M.D. performed avidity ELISAs. J.L.K. performed correlation and regression analysis. M.G., H.K., H.M.S., V.C., M.M.D., J.L.K., and J.M.F. analyzed data. M.G., J.L.K., and J.M.F. visualized data. L.E., A.P.R., and A.R.M. acquired the patient samples. A.C. provided the patient samples. A.C. and J.M.F. acquired funding. A.C. and J.M.F. supervised the study. M.G. and J.M.F. wrote the original draft. All authors reviewed and edited the manuscript.

## Acknowledgements

We thank all the study participants for their invaluable contribution. We are grateful to the clinical staff and the research coordinators for their assistance with the participant recruitment and sample collection.

## References

1. Duncan S, Eppes S. Emerging Autochthonous Transmission of Travel-Associated Vector-Borne Infections in the Continental United States. Dela J Public Health. 2025;11(1):68–71.

2. de Souza WM, Ribeiro GS, de Lima STS, de Jesus R, Moreira FRR, Whittaker C, et al. Chikungunya: a decade of burden in the Americas. Lancet Reg Health Am. 2024;30:100673.

3. CDC. Chikungunya in the United States https://www.cdc.gov/chikungunya/data-maps/chikungunya-us.html2026 [

4. O’Driscoll M, Salje H, Chang AY, Watson H. Arthralgia resolution rate following chikungunya virus infection. Int J Infect Dis. 2021;112:1–7.

5. Pedi VD, de Franca GVA, Rodrigues VB, Duailibe FT, Santos MTP, de Oliveira MRF. Burden of Chikungunya Fever and Its Economic and Social Impacts Worldwide: A Systematic Review. Trop Med Int Health. 2025;30(9):865–92.

6. Costa LB, Barreto FKA, Barreto MCA, Santos T, Andrade MMO, Farias L, et al. Epidemiology and Economic Burden of Chikungunya: A Systematic Literature Review. Trop Med Infect Dis. 2023;8(6).

7. Ng WH, Amaral K, Javelle E, Mahalingam S. Chronic chikungunya disease (CCD) clinical insights, immunopathogenesis and therapeutic perspectives.pdf. QJM: An International Journal of Medicine. 2024.

8. McCarthy MK, Morrison TE. Chronic chikungunya virus musculoskeletal disease: what are the underlying mechanisms? Future Microbiol. 2016;11(3):331–4.

9. Labadie K, Larcher T, Joubert C, Mannioui A, Delache B, Brochard P, et al. Chikungunya disease in nonhuman primates involves long-term viral persistence in macrophages. J Clin Invest. 2010;120(3):894–906.

10. Young AR, Locke MC, Cook LE, Hiller BE, Zhang R, Hedberg ML, et al. Dermal and muscle fibroblasts and skeletal myofibers survive chikungunya virus infection and harbor persistent RNA. PLoS Pathog. 2019;15(8):e1007993.

11. Hoarau JJ, Jaffar Bandjee MC, Krejbich Trotot P, Das T, Li-Pat-Yuen G, Dassa B, et al. Persistent chronic inflammation and infection by Chikungunya arthritogenic alphavirus in spite of a robust host immune response. J Immunol. 2010;184(10):5914–27.

12. Zarrella KM, Sheridan RM, Ware BC, Davenport BJ, da Silva MOL, Vyshenska D, et al. Chikungunya virus persists in joint-associated macrophages and promotes chronic disease in mice. Nat Microbiol. 2026;11(5):1302–17.

13. Chang AY, Martins KAO, Encinales L, Reid SP, Acuna M, Encinales C, et al. Chikungunya Arthritis Mechanisms in the Americas: A Cross-Sectional Analysis of Chikungunya Arthritis Patients Twenty-Two Months After Infection Demonstrating No Detectable Viral Persistence in Synovial Fluid. Arthritis Rheumatol. 2018;70(4):585–93.

14. Lum FM, Teo TH, Lee WW, Kam YW, Renia L, Ng LF. An essential role of antibodies in the control of Chikungunya virus infection. J Immunol. 2013;190(12):6295–302.

15. Smith SA, Silva LA, Fox JM, Flyak AI, Kose N, Sapparapu G, et al. Isolation and Characterization of Broad and Ultrapotent Human Monoclonal Antibodies with Therapeutic Activity against Chikungunya Virus. Cell Host Microbe. 2015;18(1):86–95.

16. Pal P, Dowd KA, Brien JD, Edeling MA, Gorlatov S, Johnson S, et al. Development of a highly protective combination monoclonal antibody therapy against Chikungunya virus. PLoS Pathog. 2013;9(4):e1003312.

17. Fox JM, Long F, Edeling MA, Lin H, van Duijl-Richter MKS, Fong RH, et al. Broadly Neutralizing Alphavirus Antibodies Bind an Epitope on E2 and Inhibit Entry and Egress. Cell. 2015;163(5):1095–107.

18. Raju S, Adams LJ, Earnest JT, Warfield K, Vang L, Crowe JE, Jr., et al. A chikungunya virus-like particle vaccine induces broadly neutralizing and protective antibodies against alphaviruses in humans. Sci Transl Med. 2023;15(696):eade8273.

19. Malonis RJ, Earnest JT, Kim AS, Angeliadis M, Holtsberg FW, Aman MJ, et al. Near-germline human monoclonal antibodies neutralize and protect against multiple arthritogenic alphaviruses. Proc Natl Acad Sci U S A. 2021;118(37).

20. Jin J, Simmons G. Antiviral Functions of Monoclonal Antibodies against Chikungunya Virus. Viruses. 2019;11(4).

21. Julie M Fox, Vicky Roy, Bronwyn M Gunn, Ling Huang, Melissa A Edeling, Matthias Mack, et al. Optimal therapeutic activity of monoclonal antibodies against chikungunya virus requires Fc-FcγR interaction on monocytes.pdf. Sci Immunol. 2019.

22. Kam YW, Lee WW, Simarmata D, Harjanto S, Teng TS, Tolou H, et al. Longitudinal analysis of the human antibody response to Chikungunya virus infection: implications for serodiagnosis and vaccine development. J Virol. 2012;86(23):13005–15.

23. Anfasa F, Lim SM, Fekken S, Wever R, Osterhaus A, Martina BEE. Characterization of antibody response in patients with acute and chronic chikungunya virus disease. J Clin Virol. 2019;117:68–72.

24. Nayak K, Jain V, Kaur M, Khan N, Gottimukkala K, Aggarwal C, et al. Antibody response patterns in chikungunya febrile phase predict protection versus progression to chronic arthritis. JCI Insight. 2020;5(7).

25. Chang AY, Encinales L, Porras A, Pacheco N, Reid SP, Martins KAO, et al. Frequency of Chronic Joint Pain Following Chikungunya Virus Infection: A Colombian Cohort Study. Arthritis Rheumatol. 2018;70(4):578–84.

26. Chang AY, Tritsch S, Reid SP, Martins K, Encinales L, Pacheco N, et al. The Cytokine Profile in Acute Chikungunya Infection is Predictive of Chronic Arthritis 20 Months Post Infection. Diseases. 2018;6(4).

27. Reinig S, Shih SR. Non-neutralizing functions in anti-SARS-CoV-2 IgG antibodies. Biomed J. 2024;47(1):100666.

28. Vidarsson G, Dekkers G, Rispens T. IgG subclasses and allotypes: from structure to effector functions. Front Immunol. 2014;5:520.

29. French MA, Tjiam MC, Abudulai LN, Fernandez S. Antiviral Functions of Human Immunodeficiency Virus Type 1 (HIV-1)-Specific IgG Antibodies: Effects of Antiretroviral Therapy and Implications for Therapeutic HIV-1 Vaccine Design. Front Immunol. 2017;8:780.

30. Martins KA, Gregory MK, Valdez SM, Sprague TR, Encinales L, Pacheco N, et al. Neutralizing Antibodies from Convalescent Chikungunya Virus Patients Can Cross-Neutralize Mayaro and Una Viruses. Am J Trop Med Hyg. 2019;100(6):1541–4.

31. Kim AS, Kafai NM, Winkler ES, Gilliland TC, Jr., Cottle EL, Earnest JT, et al. Pan-protective anti-alphavirus human antibodies target a conserved E1 protein epitope. Cell. 2021;184(17):4414–29 e19.

32. Gil-Mora J, Acevedo-Gutierrez LY, Betancourt-Ruiz PL, Martinez-Diaz HC, Fernandez D, Bopp NE, et al. Arbovirus Antibody Seroprevalence in the Human Population from Cauca, Colombia. Am J Trop Med Hyg. 2022;107(6):1218–25.

33. Ma H, Kim AS, Kafai NM, Earnest JT, Shah AP, Case JB, et al. LDLRAD3 is a receptor for Venezuelan equine encephalitis virus. Nature. 2020;588(7837):308–14.

34. Strauss JH, Strauss EG. The alphaviruses. gene expression, replication, and evolution.pdf. Microbiol Rev. 1994.

35. Zhou QF, Fox JM, Earnest JT, Ng TS, Kim AS, Fibriansah G, et al. Structural basis of Chikungunya virus inhibition by monoclonal antibodies. Proc Natl Acad Sci U S A. 2020;117(44):27637–45.

36. Fox JM, Roy V, Gunn BM, Huang L, Edeling MA, Mack M, et al. Optimal therapeutic activity of monoclonal antibodies against chikungunya virus requires Fc-FcgammaR interaction on monocytes. Sci Immunol. 2019;4(32).

37. Jin J, Galaz-Montoya JG, Sherman MB, Sun SY, Goldsmith CS, O’Toole ET, et al. Neutralizing Antibodies Inhibit Chikungunya Virus Budding at the Plasma Membrane. Cell Host Microbe. 2018;24(3):417–28 e5.

38. Chmielewski D, Schmid MF, Simmons G, Jin J, Chiu W. Chikungunya virus assembly and budding visualized in situ using cryogenic electron tomography. Nat Microbiol. 2022;7(8):1270–9.

39. Zhang J, Li C, Wu Y, Wang L, Yu J, Wang A, et al. Fc effector functions in RNA viral infections: mechanisms of antiviral immunity and implications for vaccine design. Front Immunol. 2026;17:1772257.

40. Gogesch P, Dudek S, van Zandbergen G, Waibler Z, Anzaghe M. The Role of Fc Receptors on the Effectiveness of Therapeutic Monoclonal Antibodies. Int J Mol Sci. 2021;22(16).

41. Fox JM, Roy V, Gunn BM, Bolton GR, Fremont DH, Alter G, et al. Enhancing the therapeutic activity of hyperimmune IgG against chikungunya virus using FcgammaRIIIa affinity chromatography. Front Immunol. 2023;14:1153108.

42. Jayaraman A, Walachowski S, Bosmann M. The complement system: A key player in the host response to infections. Eur J Immunol. 2024;54(11):e2350814.

43. Patil HP, Gosavi M, Kulkarni R, Mishra AC, Arankalle VA. Immunoglobulin G Subclass Response After Chikungunya Virus Infection. Viral Immunol. 2022;35(6):437–42.

44. Kam YW, Simarmata D, Chow A, Her Z, Teng TS, Ong EK, et al. Early appearance of neutralizing immunoglobulin G3 antibodies is associated with chikungunya virus clearance and long-term clinical protection. J Infect Dis. 2012;205(7):1147–54.

45. Nimmerjahn F, Ravetch JV. Fcgamma receptors as regulators of immune responses. Nat Rev Immunol. 2008;8(1):34–47.

46. C E Siegert, M D Kazatchkine, A Sjöholm, R Würzner, M Loos, Daha MR. Autoantibodies against C1q view on clinical relevance and pathogenic roles.pdf. Clin Exp Immunol. 1999.

47. Morrison TE, Fraser RJ, Smith PN, Mahalingam S, Heise MT. Complement contributes to inflammatory tissue destruction in a mouse model of Ross River virus-induced disease. J Virol. 2007;81(10):5132–43.

48. Janzic L, Kouter K. Complement and inflammasome crosstalk in chronic inflammation. Front Immunol. 2026;17:1778759.

49. Gois BM, Peixoto RF, Guerra-Gomes IC, Palmeira PHS, Dias CNS, Araujo JMG, et al. Regulatory T cells in acute and chronic human Chikungunya infection. Microbes Infect. 2022;24(3):104927.

50. Agarwal R, Chang J, Cortes FH, Ha C, Villalpando J, Castillo IN, et al. Chikungunya virus-specific CD4(+) T cells are associated with chronic chikungunya viral arthritic disease in humans. Cell Rep Med. 2025;6(5):102134.

51. Bartholomeeusen K, Affaticati F, Willems E, Dhondt E, Bartholomeus E, Maestri A, et al. Identification of a TCR signature in peripheral blood derived CD4+ T cells, associated with chronic chikungunya disease, suggests a conducive, female-biased, background immune profile. Front Immunol. 2026;17:1739100.

52. Clarke EV, Weist BM, Walsh CM, Tenner AJ. Complement protein C1q bound to apoptotic cells suppresses human macrophage and dendritic cell-mediated Th17 and Th1 T cell subset proliferation. J Leukoc Biol. 2015;97(1):147–60.

53. Teh BK, Yeo JG, Chern LM, Lu J. C1q regulation of dendritic cell development from monocytes with distinct cytokine production and T cell stimulation. Mol Immunol. 2011;48(9-10):1128–38.

54. Castellano G, Woltman AM, Schlagwein N, Xu W, Schena FP, Daha MR, et al. Immune modulation of human dendritic cells by complement. Eur J Immunol. 2007;37(10):2803–11.

55. Dias AG, Jr., Atyeo C, Loos C, Montoya M, Roy V, Bos S, et al. Antibody Fc characteristics and effector functions correlate with protection from symptomatic dengue virus type 3 infection. Sci Transl Med. 2022;14(651):eabm3151.

56. Zhang A, Stacey HD, D’Agostino MR, Tugg Y, Marzok A, Miller MS. Beyond neutralization: Fc-dependent antibody effector functions in SARS-CoV-2 infection. Nat Rev Immunol. 2023;23(6):381–96.

57. Bengtsson KL, Song H, Stertman L, Liu Y, Flyer DC, Massare MJ, et al. Matrix-M adjuvant enhances antibody, cellular and protective immune responses of a Zaire Ebola/Makona virus glycoprotein (GP) nanoparticle vaccine in mice. Vaccine. 2016;34(16):1927–35.

58. Reed SG, Tomai M, Gale MJ, Jr. New horizons in adjuvants for vaccine development. Curr Opin Immunol. 2020;65:97–101.

59. Boes M. Role of natural and immune IgM antibodies in immune responses: Elsevier; 2000. 1141–9 p.

60. Ashbrook AW, Burrack KS, Silva LA, Montgomery SA, Heise MT, Morrison TE, et al. Residue 82 of the Chikungunya virus E2 attachment protein modulates viral dissemination and arthritis in mice. J Virol. 2014;88(21):12180–92.

