## Supplemental Figures for "Enhanced early IgG-mediated complement deposition in the development of chronic chikungunya virus disease"

**Supplemental Figures and Table**

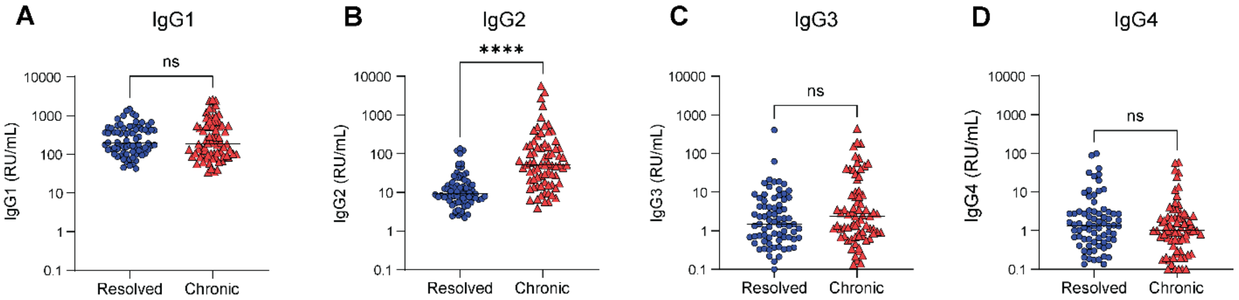


**Fig S1. Related to Figure 1. Increased bulk IgG2 in chronic samples.** Bulk levels of (**A**) IgG1, (**B**) IgG2, (**C**) IgG3, and (**D**) IgG4 in the purified IgG from the down-selected samples were quantified by ELISA compared to a standard curve. Each dot represents one individual, and the bars indicate the median. RU stands for relative units. Statistical analysis was performed using a Mann-Whitney test; ***p < 0.0001, ns = not significant.

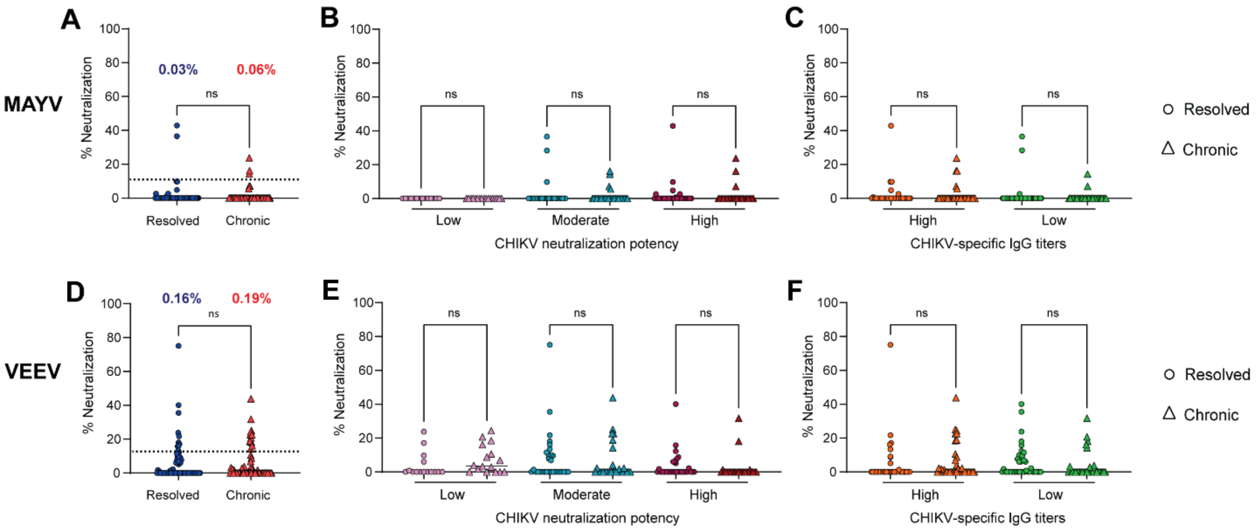


**Fig S2. Related to Figure 1. No difference in MAYV or VEEV neutralization between resolved and chronic groups.** Purified IgG from the down-selected samples was evaluated for (**A-C**) MAYV or (**D-F**) VEEV neutralization by FRNT. IgG samples were considered positive for neutralization if ≥10% neutralization was achieved at 10 μg/ml. The percent of samples positive for neutralization against (**A**) MAYV or (**D**) VEEV. The percent neutralization of MAYV or VEEV was separated by (**B** and **E**) CHIKV neutralization or (**C** and **F**) CHIKV-specific IgG in resolved and chronic samples. Each dot represents one individual. Bars indicate the median. The dotted line indicates 10% neutralization. Statistical comparisons were performed using a Mann-Whitney test.

**Table S1. Related to Figure 1. Sample down-selection groups**

| CHIKV neutralization potency groups | $IC_{50}$ value (µg/ml) | Resolved | | | Chronic | | | Total | | |
|---|---|---|---|---|---|---|---|---|---|---|
| | | N | % Female | Age[1] | N | % Female | Age | N | % Female | Age |
| Low | > 8 | 14 | 57.1 | 50.2 (18-68) | 16 | 87.5 | 47.4 (25-64) | 30 | 72.3 | 48.8 (18-68) |
| Moderate | 2 – 8 | 30 | 83.3 | 49.0 (21-84) | 32 | 78.1 | 48.5 (21-88) | 62 | 80.7 | 48.7 (21-88) |
| High | < 2 | 30 | 90.0 | 50.6 (14-90) | 24 | 91.6 | 48.6 (30-80) | 54 | 90.8 | 49.6 (14-90) |

| CHIKV-specific IgG titer groups | IgG endpoint dilution | Resolved | | | Chronic | | | Total | | |
|---|---|---|---|---|---|---|---|---|---|---|
| | | N | % Female | Age[1] | N | % Female | Age | N | % Female | Age |
| High | 1600 – 6400 | 33 | 90.9 | 48.8 (18-90) | 40 | 87.5 | 48.7 (21-88) | 73 | 89.2 | 48.7 (18-90) |
| Low | 400 – 800 | 41 | 73.0 | 49.1 (14-77) | 32 | 81.25 | 48.6 (17-80) | 73 | 77.2 | 48.9 (14-80) |

[1]Median age is shown with range
